# Intense and sustained pain reduces cortical responses to auditory stimuli: implications for the interpretation of Heterotopic Noxious Conditioning Stimulation in humans

**DOI:** 10.1101/460576

**Authors:** DM Torta, FA Jure, OK Andersen, JA Biurrun Manresa

**Author notes:** Corresponding author: Dr. Diana Torta, Research group of Health Psychology, Katholieke Universiteit Leuven Tiensestraat 102, 3000 Leuven Belgium; Institute of Neuroscience, Université catholique de Louvain Avenue Mounier 53, 1200 Brussels Belgium.

## Abstract

Phasic pain stimuli are inhibited when they are applied concomitantly with a conditioning tonic stimulus at another body location (Heterotopic Noxious Conditioning Stimulation, HNCS). While this effect is thought to rely on a spino-bulbo-spinal mechanism in animals (Diffuse Noxious Inhibitory Controls, DNIC), the underlying neurophysiology in humans may further involve other pathways. In this study, we investigated the role of supraspinal mechanisms in HNCS by presenting auditory stimuli during a conditioning tonic painful stimulus (the Cold Pressor Test, CPT). Considering that auditory stimuli are not conveyed through the spinal cord, any changes in brain responses to auditory stimuli *during* HNCS can be ascribed entirely to supraspinal mechanisms. High-density electroencephalography (EEG) was recorded during HNCS and auditory stimuli were administered in three blocks, *before*, *during*, and *after* HNCS. Nociceptive Withdrawal Reflexes (NWRs) were recorded at the same time points to investigate spinal processing. Our results showed that AEPs were significantly reduced *during* HNCS. Moreover, the amplitude of the NWR was significantly diminished *during* HNCS in most participants. Given that spinal and supraspinal mechanisms operate concomitantly during HNCS, the possibility of isolating their individual contributions to DNIC-like effects in humans is questionable. We conclude that the net effects of HCNS cannot be measured independently from attentional/cognitive influences.

## Introduction

Pain is modulated by inhibitory and facilitatory mechanisms at both spinal and supraspinal levels (Millan, 2002; Wiech, 2016). In animals, the responses of convergent wide dynamic range neurons (WDR) of the dorsal horn are strongly inhibited when a nociceptive stimulus is applied outside of their excitatory receptive field (Le Bars, 2002). This effect has been termed Diffuse Noxious Inhibitory Control (DNIC) and is thought to rely on a spinal-bulbar-spinal loop (Le Bars et al., 1979a; b; Willer et al., 1984; De Broucker et al., 1990; Willer et al., 1990; Le Bars et al., 1992; Villanueva & Le Bars, 1995). In humans, the cortical and spinal responses to a noxious stimulus can also be modulated by another noxious stimulus applied at a remote body part, a phenomenon termed ‘Heterotopic Noxious Conditioning Stimulation’, HNCS (Le Bars et al., 1979a; b; Villanueva & Le Bars, 1995; Pud et al., 2009; Kennedy et al., 2016). This similarity led to the suggestion that the same mechanisms act in humans and in animals (Kennedy et al., 2016; Bannister & Dickenson, 2017). Nevertheless, studies have proposed that the reduction in pain perception and nociceptive responses in humans, measured by functional magnetic resonance imaging (fMRI) and event-related potentials (ERPs), can be due to mechanisms different from the spino-bulbar ones, i.e. different from the DNIC effects Sprenger et al., 2011; Torta et al., 2015). Indeed, HNCS reduces somatosensory evoked potentials (SEPs) elicited by stimuli that activate Aβ-fibers Torta et al., 2015; Rustamov et al., 2016). Considering that SEPs are thought to reflect the engagement of fibers conveyed in the dorsal columns and not in the human spino-thalamic tract (Kakigi et al., 1991; Treede et al., 1991; Iannetti et al., 2001), their reduction during HNCS would advocate against modulation by DNIC mechanisms (Torta et al., 2015).

Nevertheless, an indirect pathway was observed in animals between the dorsal column and the WDR neurons that could at least partially contribute to the reduction in brain responses to non-nociceptive somatosensory stimuli (Hazlett et al., 1972). In this study we evaluated the effects of HNCS on responses to auditory stimuli. We recorded auditory evoked potentials (AEPs) before, during and after HNCS, and we also recorded the NWR at the same time points, as a control measure for spinal responses to noxious stimuli. The working hypothesis was that a reduction in AEPs during HNCS would necessarily mean that concomitant mechanisms originating at the supraspinal level are recruited.

## Methods

### Participants

The experiment was conducted on 16 healthy volunteers (7 women and 9 men, aged 20 to 35 years) who had no history of neurological, psychiatric, dermatological or chronic pain disorders, and no recent history of psychotropic or analgesic drug use. The protocol was approved by the Region Nordjylland (Denmark) ethics committee (VN-20150038). Written informed consent was obtained from all participants.

### Experimental design

At the beginning of the session, volunteers were given a detailed explanation of the experimental procedures and were familiarized with the experimental setup and task, the auditory and noxious stimuli and the rating procedures. Participants were comfortably seated in a supine position on an adjustable bed, with back support at 120° and knees flexed at 30°, relative to the horizontal.

The experiment consisted of three blocks: *before*, *during* and *after* HNCS. In each block, participants received auditory and electrical stimuli. Electrical stimuli were used to elicit the NWR (see following paragraphs for details on how NWRs were obtained). The two types of stimuli (auditory and electrical) were presented in different blocks in a counterbalanced order across participants. Upon experimenter’s request, participants had to rate the perceived intensity of auditory and electrical stimuli on a Numerical Rating Scale (NRS) ranging from 0 to 10 (0 = did not feel anything, 10 = the maximal tolerable intensity possible for this kind of stimulus). The ratings were asked for five auditory stimuli and three electrical ones. This procedure was preferred to an evaluation after each stimulus as it required a stable level of attention to the stimuli (to comply with the random request) at the same time minimizing artifacts in the electroencephalogram (EEG) related to muscular activity.

### Stimuli

#### Auditory stimuli

A total of 20 brief computer-controlled stimuli were presented via two loudspeakers (Altec Lansing Technologies, Inc.; Milford, PA 18337, USA) placed behind the participant and aligned with their body midline. The same intensity was used for all participants. Each stimulus consisted of a pure sinusoidal signal with a 3- ms period, lasted 100 ms, and measured approximately 80 dB. The inter-stimulus interval (ISI) ranged between 5 and 8 s.

#### Electrical stimuli

A total of 10 stimuli were delivered through a self-adhesive surface electrode (type 700, 20 x 15 mm, Ambu A/S, Denmark) placed on the arch of the foot, acting as cathode. The anode electrode (50 x 90 mm, Pals, Axelgaard Ltd., Fallbrook, California, USA) was placed on the dorsum of the foot. This procedure ensured the activation of nociceptors at the arch of the foot (Frahm et al., 2013). Stimulation intensity was twice the reflex threshold (RTh), assessed on the biceps femoris (BF) muscle. Each stimulus consisted of five constant-current, 1-ms rectangular pulses at 200 Hz (Noxitest IES 230, Aalborg University, Denmark), felt as a single pricking stimulus. The ISIs ranged from 8 to 10s.

#### Heterotopic noxious conditioning stimulation (HNCS)

In each of the three blocks, participants had to immerse their right hand in a bath with circulating water. In the *before* and *after* blocks, the temperature of the water was lukewarm, perceived as neither cold nor hot by the participants. The temperature was recorded at the beginning and at the end of each block (*before* block: temperature at the beginning 30.6 ± 0.8 °C, at the end 30.4 ± 0.8 °C; *after* block: temperature at the beginning 30.16 ± 0.8 °C, at the end 30.4 ± 0.7 °C). In the HNCS session, the temperature of the circulating water was below 3 °C (2.6 ± 0.4 °C at the beginning and 2.9 ± 0.4 °C at the end). In each block, recordings began 10 s after immersing the hand in the water bath. A five- to seven-minute rest period separated the blocks depending of the hand temperature. The exact timing between the *during* and *after* blocks was determined on the time needed for the immersed hand to return to baseline temperature (see Figure 1 for details of the experimental setup).

**Figure 1.**
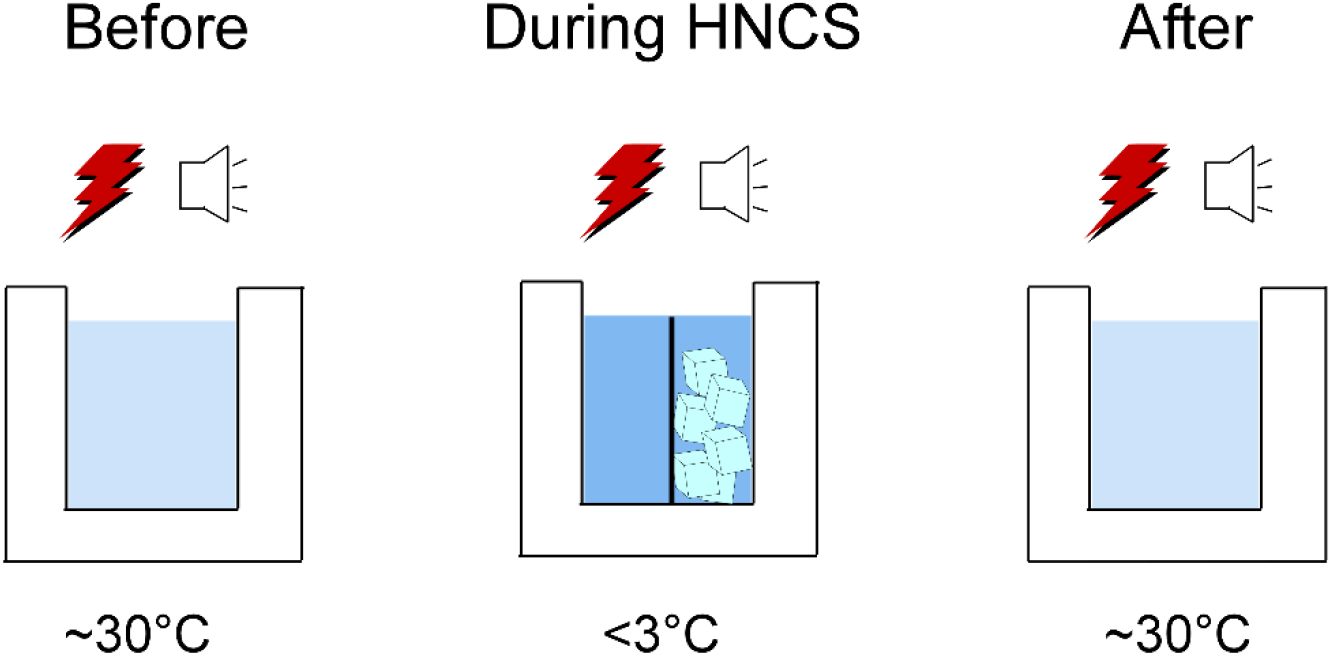
AEPs and NWRs were recorded in separate blocks, before, during and after HNCS. The temperature of the water was around 30 degrees in the before and after session and was kept below 3 degrees in the during session by means of an electrical pump. Twenty auditory stimuli and 10 electrical stimuli were used to evoke AEPs and NWRs.

**Figure 2.**
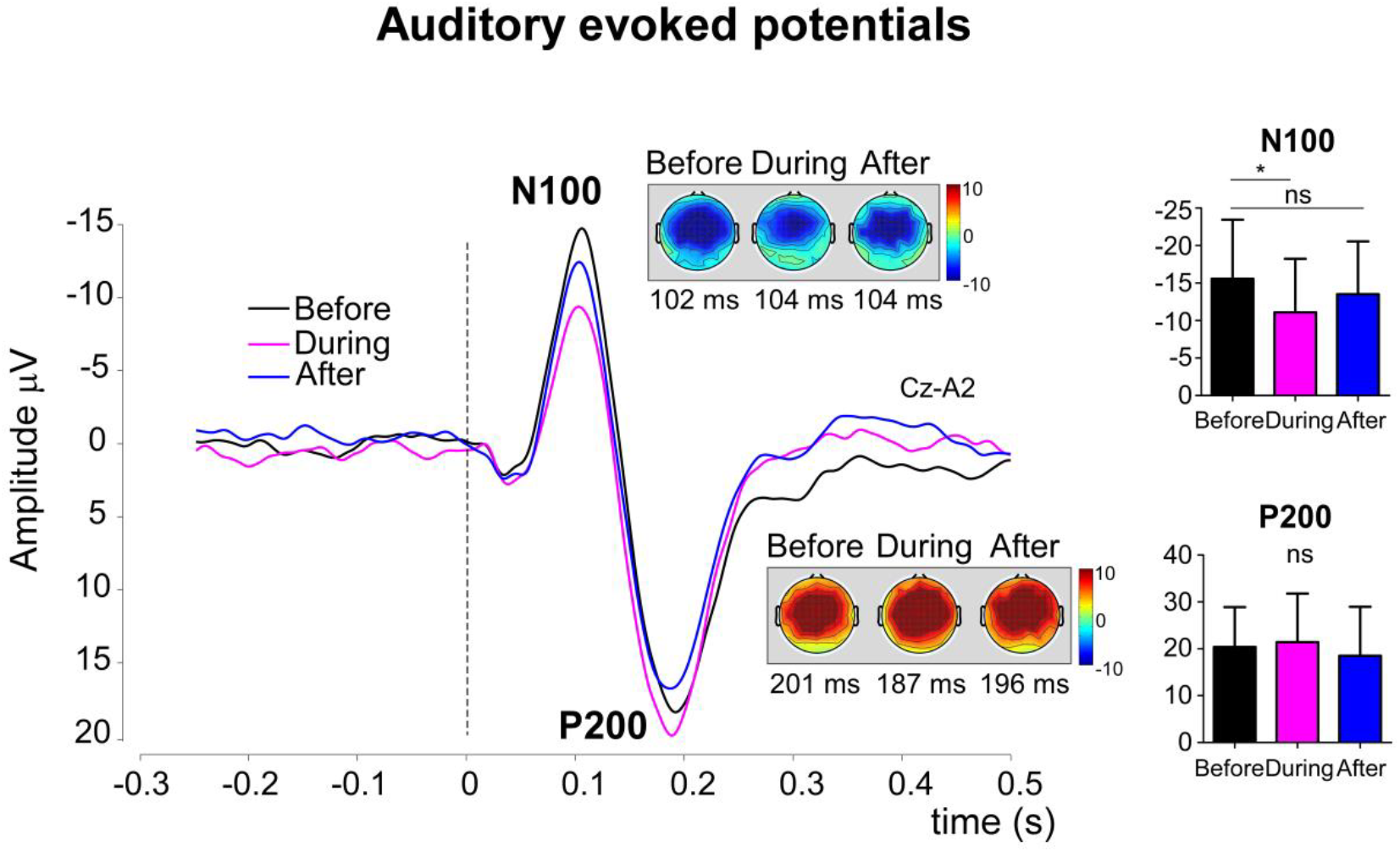
Auditory evoked potentials (AEPs). The magnitude of the N100 was reduced in the during session as compared to before, but no differences emerged between the after and before blocks. Notably, the amplitude of the response increased significantly in the after block (p = 0.03), indicating that the reduction was not only due to habituation. No significant effects were observed in the amplitude of the P200. Bars represent standard deviations (N = 15).

### Electromyographic recordings (EMG)

EMG was recorded using surface electrodes (type 720, Ambu A/S, Denmark) placed with an inter-electrode distance of 20 mm over the belly of Biceps Femoris (BF) muscle along the main direction of the muscle fibers. Before the placement of the electrodes, the skin was lightly abraded and cleaned to decrease the electrode impedance. EMG signals were sampled at 2400 Hz, amplified (up to 20000 times), band-pass filtered (5-500 Hz), displayed and stored between 500 ms before and 2000 ms after stimulation onset.

### Nociceptive Withdrawal Reflexes (NWRs): threshold and detection

An automated standardized staircase procedure was used to determine the Reflex Threshold (RTh). The RTh was identified by first applying an ascending staircase using steps of 2 mA until a NWR was detected in the BF muscle (see NWR detection criteria below). Then, a decrement of the intensity was applied in steps of 1 mA until no NWR was detected. Subsequently, three ascending and three descending staircases were applied. The RTh was defined as the average intensity of the last three two peaks and troughs (Jensen et al., 2015b).

The NWR was detected using a peak z-score criterion. The interval peak z-score was calculated as the difference between the peak value in the reflex quantification interval (60-180 ms post-stimulation) and baseline mean value, divided by the standard deviation of the pre-stimulation background activity. A NWR was detected when the interval peak z-score was larger than 12 (Rhudy & France, 2007).

To assess the conditioning effect, the NWR size was quantified. The root-mean-square (RMS) was used to calculate the NWR sizes in the reflex window interval (see above). The RMS of all trials was calculated and averaged per block and participant.

### Electroencephalographic recordings (EEG)

EEG was recorded continuously at a 2400 Hz sampling rate with a 60-channel g.tec© system. Data was recorded using the right earlobe (A2) as reference and bandpass filtered between 0.5-30 Hz. Off line analyses were performed using Letswave 6 (http://www.nocions.org/letswave). The signal was then segmented into epochs ranging from 500 ms before to 500 ms after stimulus onset. Artifact related to eye-blinks (e.g. ICs containing a large frontal distribution) and/or slow drifts were removed using an Independent Component Analysis (ICA) based on a RUNICA unconstrained algorithm (Jung et al., 2000) The epochs were baseline corrected by subtracting the mean amplitude of the signal in the -500 to 0 ms interval. Epochs exceeding ±100 μV were rejected as considered contaminated by artifacts.

Artifact-free responses were averaged for each participant and condition yielding three grand-averages corresponding to the *before, during* and *after* blocks. ERPs elicited by auditory stimuli (Auditory Evoked Potentials, AEPs) were measured at the vertex electrode (Cz), to allow a comparison with previous studies (Plaghki et al., 1994; Torta et al., 2015; Rustamov et al., 2016). The P50 was identified as the most positive deflection occurring between 25 and 70 ms. The N100 was defined as the most negative deflection occurring in the interval between 80 and 180 ms post stimulus; the P200 as the most positive deflation following the first negative wave and appearing in the time window between 200 and 300 ms. Components were clearly identified in all participants.

### Statistics

Data were analyzed using IBM SPSS Statistics 19 (IBM Corp. Released 2010. IBM SPSS Statistics for Windows, Version 19.0. Armonk, NY: IBM Corp). Repeated-measures analysis of variance (RM-ANOVA) with the factor ‘Block’ (three levels, *before*, *during* and *after*) were performed, and the Greenhouse-Geisser correction was used when the sphericity assumption was not met, as assessed by the Mauchly test. Post hoc t tests were carried out in case of significant effects. Non-parametric Friedman ANOVA was performed to assess changes in NWR amplitudes, since data were non-normally distributed (assessed with the Shapiro-Wilk test). Post hoc Wilcoxon tests were carried out in case of significant effects. The critical alpha level was set at p = 0.05 and corrected for multiple comparisons using the Bonferroni procedure.

## Results

### Auditory evoked potentials

One of the participants had to be excluded due to large electrical artifacts at the Cz electrode. Therefore, the final sample for the EEG analysis consisted of 15 subjects. The amplitude of the N100 was reduced *during* HNCS (p = 0.003) but did not show any significant difference compared to baseline levels in the *after* block. Notably, also the comparison *during* vs. *after* resulted in a significant difference (p = 0.03). No effects were observed on the amplitude of the P50 the P200, and the latency of all measured components.

### NWR

The NWR sizes did not show significant differences between blocks (see Figure 3).

Four (out of the sixteen) participants were considered ‘Non-responders’ in view of an enhancement of the NWR size in the *during* block. When the analyses were performed without these participants, a significant difference of the NWR size was observed across blocks Χ^2^ = 11.167, p = 0.0037. The NWR was significantly reduced *during* HNCS (Z = -3.059, *p* = 0.004). In contrast, no significant difference was found when comparing the *before* and *after* blocks. These results can be found in Supplementary Figure 1.

**Figure 3.**
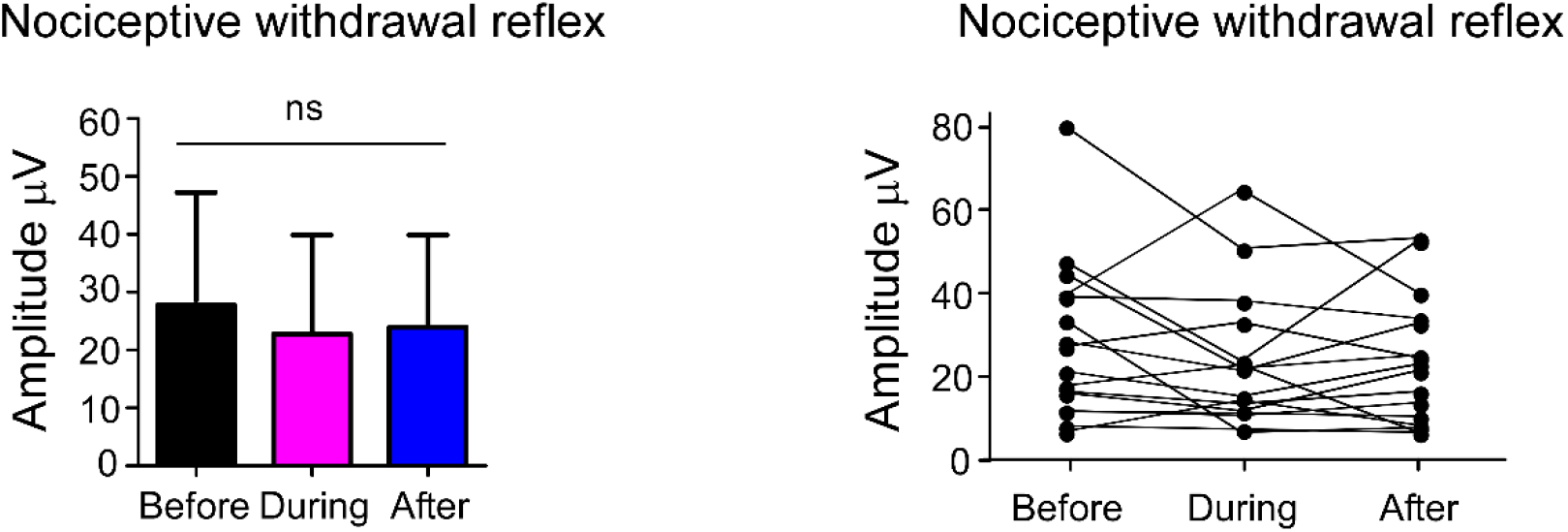
The magnitude of the nociceptive withdrawal reflex did not change significantly across blocks when all participants (N=16) were considered.

### Ratings

The perceived intensity of auditory stimuli remained unchanged across blocks. A main effect of ‘Block’ was observed for electrical stimuli (F_G-G_(1.4,19.6) = 7.759, p = 0.007, partial η^2^ = 0.357). The stimuli were not perceived as less intense *during* HNCS but felt more intense *after* HCNS (p = 0.013) (see Figure 4).

**Figure 4.**
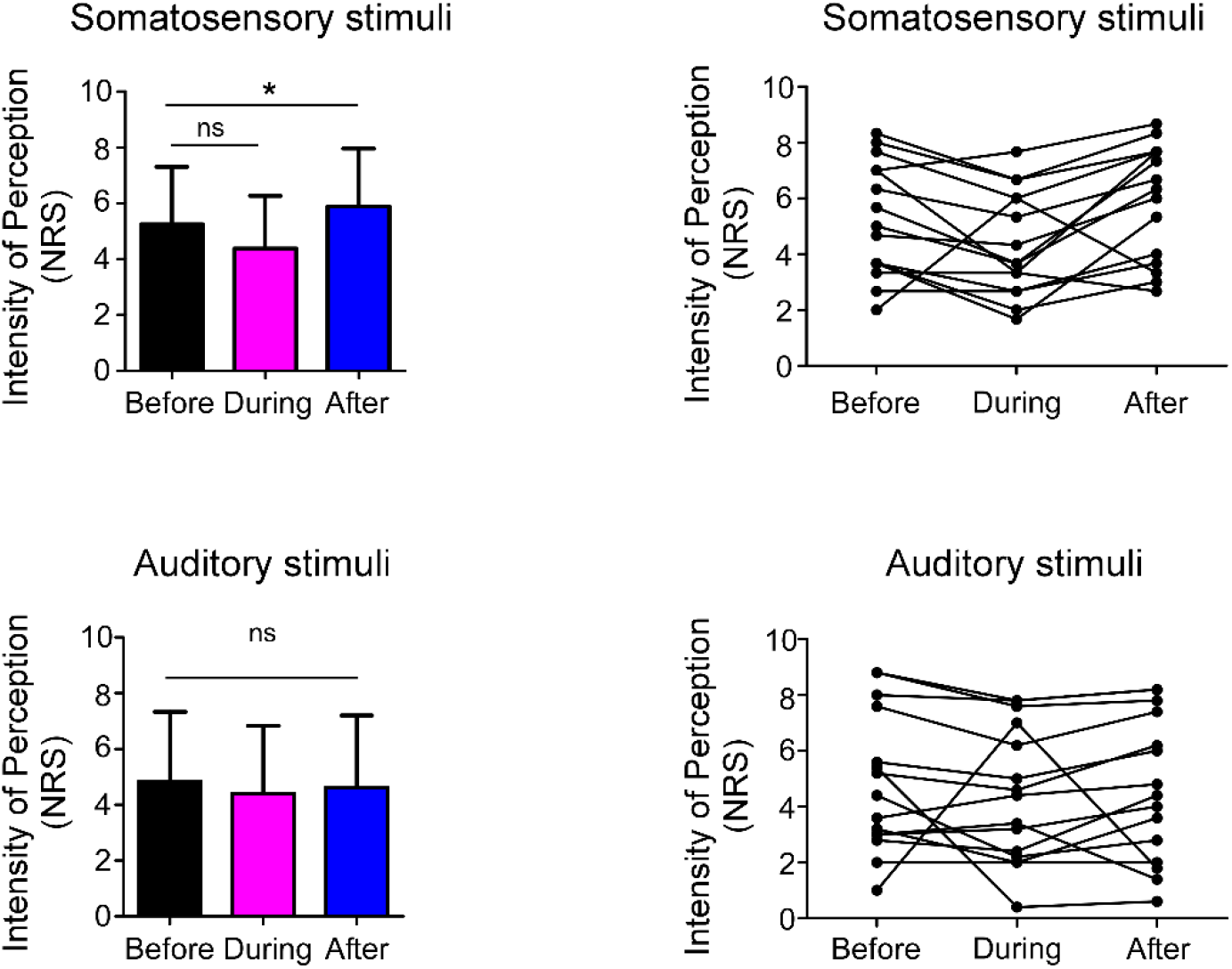
Left columns: average and standard deviation of the perceived intensity of the stimuli at the group level. Individual level data are provided in the right-hand columns. Bars represent standard deviations (N = 16).

## Discussion

This study addressed whether it is possible to consider spinal-bulbo-spinal mechanisms as operating independently from supraspinal, possibly attentional, mechanisms during HNCS. The reduction in AEPs clearly indicates that during HNCS supraspinal inhibitory mechanisms are recruited. Therefore, our results demonstrate that several mechanisms act concomitantly during HNCS and the relative contribution of each of these mechanisms cannot be readily quantified. Indeed, the supraspinal mechanisms acting over the AEPs may be involved also in the reduction of spinal responses, given that an attentional modulation of the NWR has been observed (Bjerre et al., 2011). Furthermore, a reduction in AEPs during HNCS was observed also in those participants who showed larger NWRs during HNCS, suggesting that competing spinal/supraspinal mechanisms can be concomitantly acting.

### Supraspinal processes during HNCS

The magnitude of the vertex negative component of the AEPs (N100) was significantly reduced during HNCS, and returned to baseline levels in the after session, when the hand was immersed in lukewarm water. This result confirms and expands previous physiological findings showing that HNCS modulates brain responses to non-nociceptive somatosensory inputs (Torta et al., 2015). Indeed, it confirms that HNCS recruits mechanisms, possibly attentional, which act entirely at the supraspinal level, independently from spinal loops. Notably, Sprenger et al., (Sprenger et al., 2011) have reported a reduction in the BOLD signal during HNCS in multimodal areas comprising the insula, the cingulate cortex and the somatosensory cortices. Nevertheless, considering that the authors used nociceptive stimuli and that concomitant spinal and supraspinal fMRI recording is challenging, it was impossible to establish whether the observed modulation occurred entirely at the supraspinal level, or the reduction in BOLD responses was a consequence of spinal gating.

The present study offers a new interpretation to the psychophysiological effects of HNCS. In a previous report, Moont and colleagues (2010) attempted to determine whether HNCS and distraction modulate pain via the same or distinct mechanisms. The assumption was that these effects would be additive in case of independence of the mechanisms; in other words, that the effects of combined HNCS and distraction would be larger than HNCS alone. The results showed indeed that the reduction in pain ratings under the combined stimulation was significantly greater than under the conditioning stimulation alone (immersion of the hand in a hot bath). Importantly however, the reduction in the combined stimulation condition was not different from the distraction stimulation. The authors’ conclusion was that the significant additive effects of pain inhibition when distraction was combined with tonic conditioning pain supported the view that HCNS and distraction act independently. Nevertheless, for these statements to be true, the combined effect should have been the net sum of the effects of HNCS and of distraction. This was not the case, indeed, the combined effect was not different from distraction alone (Moont et al., 2010; van Wijk & Veldhuijzen, 2010), see also (Lautenbacher et al., 2007). Thus, another account is possible, namely that attentional mechanisms are per se recruited during HNCS, and they also modulate the response in a way that spinal and supraspinal components cannot be readily quantified in healthy volunteers using standard assessment procedures, given the strong attentional component inherent to the conditioning paradigm.

Notably, the major difference of this study in comparison with previous studies was in the choice of not using an attentional paradigm to modulate the effects of HNCS (e.g. (Ladouceur et al., 2012) or compare HNCS-induced analgesia to distraction-induced analgesia (Lautenbacher et al., 2007; Moont et al., 2010). Rather, the aim was to show that HNCS recruits *per se* attentional mechanisms which may be responsible for many of the previous findings (Bouhassira et al., 1992; Villanueva & Le Bars, 1995; Lapirot et al., 2009; Lapirot et al., 2011; Chebbi et al., 2014). Noteworthy, we use in this report the generic terms ‘attention’, ‘distraction’, or ‘cognition’. In reality, these concepts refer to several distinct processes, supported by different networks (for a review in the pain field see Torta et al., 2017). Considering that Moont and colleagues (2010) specifically tested for the effects of *spatial attention*, whereas in our study we did not manipulate attention directly, we can conclude that effects occur at the supraspinal level, but we cannot go as far as interpreting whether the N100 reduction is related to distraction, mental effort to cope with the intense ongoing pain or to a gating of the sensory input. All these possibilities refer to different physiological processes (e.g. Raz, 2004; Raz & Buhle, 2006; Torta et al., 2017). Certainly, a reduction in auditory responses cannot be explained by a descending modulatory inhibition involving specifically nociceptive circuits.

### HNCS effects on perception and NWR

The possibility that HNCS modulates the perceived intensity of somatosensory stimuli has received ambiguous support (Rhudy et al., 2006; Defrin et al., 2007). Recently, Torta et al., (2015) did not find evidence in that direction. Rustamov and colleagues (2016) reported instead a decrease in the perceived intensity of the electrical shocks of both lower and higher intensity *during* HNCS, but a persisting reduced perception only for stimuli of lower intensities. In contrast, the present study evidenced an increase in the perceived intensity of electrical stimuli a*fter* HNCS as compared to *before*. Considering the inconsistency of the results, and the relative paucity of data it is arduous to provide a unifying view or an explanation for such divergent findings. At present we can only speculate on the possibility that different outcomes were driven by different experimental approaches.

Previous studies have shown a depression of the NWR during HNCS (Terkelsen et al., 2001; Biurrun Manresa et al., 2011; Rustamov et al., 2016), whereas only one reported a failure to induce NWR inhibition in a specific experimental paradigm (Terkelsen et al., 2001). In this sense our results are surprising. However, differences in experimental protocols should be considered and could explain the lack of modulation at the group level in the present study. Here, we have applied electrical stimuli on the arch of the foot and not on the sural nerve, as in previous reports. Besides, the stimulation of the arch of the foot seems to be more reliable in eliciting a NWR (Jensen et al., 2015a), and limits the risk of drop-outs during the stimulation due to intolerable pain (Biurrun Manresa et al., 2014). Indeed, stimulation at the arch of the foot is usually performed at lower intensities compared to the sural nerve, resulting in smaller NWR and possibly making changes in amplitude harder to detect.

Furthermore, it was observed that 4 volunteers showed an increase, rather than a decrease of the NWR amplitude during HNCS, in line with other recent studies (e.g. (Potvin & Marchand, 2016). It is becoming increasingly accepted that some participants do not show an inhibitory effect of HNCS at the perceptual level (Youssef et al., 2016b; a), although presently there is no standardized procedure on how to report this observation (Kennedy et al., 2016). However, recent attempts to quantify the proportion of responders and non-responders based on meaningful CPM effect sizes showed that there are large variations across protocols, and large proportion (from 11 to 73%) of subjects could be potentially classified as CPM non-responders (Locke et al., 2014; Vaegter et al., 2018).

### Conclusion

As previous reports (Torta et al., 2015), we conclude that several mechanisms co-occur during HNCS in humans. However, we also suggest that this concomitant occurrence hampers the possibility of attributing any modulatory effect during HNCS to DNIC-like mechanisms alone.

**Table 1.** Summary of the results. Values in red indicate significant differences

|  | Main effect of Block | Before vs. During* | Before vs. After* |
| --- | --- | --- | --- |
| **NWR (Amplitude measured at the Biceps Femoris)** |  |  |  |
| All Subjects (n = 16) | $X^2 = 3.125$,<br>$p = 0.209$ | | |
| Responders (n = 12) | $X^2 = 11.167$,<br>$p = 0.004$ | $Z = -3.059$,<br>$p = 0.004$ | $Z = 0.627$,<br>$p > 0.999$ |
| **Perceived intensity** |  |  |  |
| Auditory stimuli | $F(2,28) = 0.382$, $p = 0.686$ | | |
| Electrical stimuli | $F(2,28) = 7.759$, $p = 0.007$ | $T(14) = 1.971$, $p = 0.069$ | $t(14) = -2.846$, $p = 0.013$ |
| **Auditory evoked potentials** |  |  |  |
| N100 amplitude | $F(2,28) = 7.016, p = 0.003$ | $t(14) = -4.218, p = 0.001$ | $t(14) = -1.441, p = 0.172$ |
| N100 latency | $F(2,28) = 0.318, p = 0.730$ | | |
| P200 amplitude | $F(2,28) = 1.294, p = 0.290$ | | |
| P200 latency | $F(2,28) = 2.566, p = 0.095$ | | |

\* the critical alpha level was set at $p=0.05$ and corrected for multiple comparisons

## Acknowledgments

DMT was supported by the Fund for Scientific Research of the French-speaking Community of Belgium (F.R.S.-FNRS) and by the Asthenes long-term structural funding-Methusalem grant (# METH/15/011) by the Flemish Government of Belgium. Other acknowledgments go to the Danish National Research Foundation (DNRF121).

## Author Contributions

DM conceived the research, DM, FAJ, JBM discussed the methods and setup, FAJ collected the data, all authors wrote the manuscript and critically reviewed it.

## Data statement

The full set of data is available upon request to the first author.

## Abbreviations

DNIC: Diffuse Inhibitory Controls
EEG: Electroencephalography
HNCS: Heterotopic Noxious Conditioning Stimulation
NWR: Nociceptive Withdrawal Response
SEP: Somatosensory Evoked Potential

